# Chromosome size affects sequence divergence between species through the interplay of recombination and selection

**DOI:** 10.1101/2021.01.15.426870

**Authors:** Anna Tigano, Ruqayya Khan, Arina D. Omer, David Weisz, Olga Dudchenko, Asha S. Multani, Sen Pathak, Richard R. Behringer, Erez L. Aiden, Heidi Fisher, Matthew D. MacManes

## Abstract

The structure of the genome shapes the distribution of genetic diversity and sequence divergence. To investigate how the relationship between chromosome size and recombination rate affects sequence divergence between species, we combined empirical analyses and evolutionary simulations. We estimated pairwise sequence divergence among 15 species from three different Mammalian clades - *Peromyscus* rodents, *Mus* mice, and great apes - from chromosome-level genome assemblies. We found a strong significant negative correlation between chromosome size and sequence divergence in all species comparisons within the *Peromyscus* and great apes clades, but not the *Mus* clade, suggesting that the dramatic chromosomal rearrangements among *Mus* species may have masked the ancestral genomic landscape of divergence in many comparisons. Our evolutionary simulations showed that the main factor determining differences in divergence among chromosomes of different size is the interplay of recombination rate and selection, with greater variation in larger populations than in smaller ones. In ancestral populations, shorter chromosomes harbor greater nucleotide diversity. As ancestral populations diverge, diversity present at the onset of the split contributes to greater sequence divergence in shorter chromosomes among daughter species. The combination of empirical data and evolutionary simulations revealed that chromosomal rearrangements, demography, and divergence times may also affect the relationship between chromosome size and divergence, and deepen our understanding of the role of genome structure on the evolution of species divergence.

## Introduction

Chromosomes are the fundamental unit of inheritance of the nuclear DNA in all eukaryotic species and their evolution goes arm in arm with organismal evolution. Not only the sequence, but also the size, shape, structure, and number of chromosomes can vary between species, populations, and even individuals within a population (Hauffe and Searle 1993; Graphodatsky et al. 2011; Dion-Côté et al. 2017; Moura et al. 2020). Chromosome evolution is therefore crucial to our understanding of the evolution and maintenance of biodiversity. Despite this, our current understanding of these relationships is limited. The rapidly increasing number of chromosome-level genome assemblies has started to shed light on the role of chromosomes in the genomic distribution of genetic diversity within and between species. For example, chromosome structure, including the location of telomeres or centromeres, can be a strong predictor of the position of dips and peaks in nucleotide diversity (*π*) and sequence divergence (*d*) within a chromosome (Butlin 2005; Smukowski and Noor 2011; Burri et al. 2015; Sardell et al. 2018; Tigano et al. 2021; Robinson et al. 2021). The heterogeneous distribution of *π* and *d* in the genome is apparent among chromosomes too (Dutoit et al. 2017; Murray et al. 2017; Henderson and Brelsford 2020; Robinson et al. 2021), but the role of genome structure in generating this distribution, including the number and size of chromosomes in a genome, is less clear.

To avoid the production of aberrant gametes, the correct segregation of chromosomes during meiosis requires that chromosomes undergo at least one cross-over per event (Mather 1938; Hassold and Hunt 2001), which results in shorter chromosomes experiencing overall proportionally higher recombination rates. In fact, a significant relationship between chromosome size and recombination rate has been reported in many species (but not all) from fungi to mammals (Kaback et al. 1992; Jensen-Seaman et al. 2004; Pessia et al. 2012; Farré et al. 2013; Kawakami et al. 2014; Haenel et al. 2018). Chromosome size is also inversely correlated with *π* in some species of birds and mammals (Dutoit et al. 2017; Murray et al. 2017; Tigano et al. 2020; Robinson et al. 2021), but not others (Pessia et al. 2012; Dutoit et al. 2017). Further, higher *d* in microchromosomes (< 20 Mb) relative to macrochromosomes (> 40 Mb) has been observed in several bird species (Delmore et al. 2018). While these studies highlight the intricate relationship between chromosome size, recombination rate, nucleotide diversity, and sequence divergence, many other factors likely contribute to this relationship. Currently, our understanding of these additional factors is limited by both theory and lack of empirical data.

Investigating the factors shaping the levels and patterns of sequence divergence between species is fundamental to understand the molecular mechanisms underlying the process of adaptation and speciation. Nonetheless, the relative contribution of recombination to divergence among species has rarely been directly investigated, especially at the chromosome scale, in species other than humans and their closest relatives, the great apes (Hellman et al. 2003; Phung et al. 2016). However, a recent study used chromosome size as a proxy for recombination rate to test how genome structure affected divergence among eight avian sister species pairs, and reported significantly higher divergence in microchromosomes than in macrochromosomes (Delmore et al. 2018), but it did not address the mechanisms underpinning this relationship.

Although the correlation between recombination rate and *π* has often been reported (Begun and Aquadro 1992; Nachman 2001; Cutter and Choi 2010), the factors and their relative roles in determining this relationship are less clear (Ellegren and Galtier 2016). Non-crossover gene conversion (gene conversion hereafter) and selection, including linked selection, are among the factors most commonly invoked to explain the correlation between recombination and *π*. Gene conversion is the process by which double-strand DNA breaks during meiosis are repaired using homologous sequence as template without crossing-over, and although it affects shorter sequences than other types of crossing-over events do, it can increase diversity and affect divergence among populations and species (Korunes and Noor 2017). Under a purely neutral model of evolution, π is determined by the effective population size (*N_e_*) and the mutation rate (*μ*), as expressed by the equation *π*=4*N_e_μ* (Tajima 1983). In this model, higher recombination rate may increase π in smaller chromosomes if recombination results in the introduction of mutations through gene conversion (Coop and Przeworski 2007; Arbeithuber et al. 2015). For example, a study on humans showed a correlation between recombination, diversity, and divergence to the chimpanzee and the baboon, and explained these relationships with a pure neutral model entailing recombination-associated variation in mutation rates (Hellmann et al. 2003). In contrast, under a non-neutral evolutionary model, selection reduces diversity in the genomic regions surrounding beneficial or deleterious mutations via selective sweeps and background selection, respectively, with a greater diversity-reducing effect in areas of low recombination (Smith and Haigh 1974; Wiehe and Stephan 1993; Hudson and Kaplan 1995), even though purifying selection can also counteract the loss of diversity due to background selection by associative overdominance if the deleterious variant is recessive (Ohta 1971; Gilbert et al. 2020). Support for the role of selection comes from another study on humans, showing how background selection in the ancestral population affects neutral divergence by reducing diversity in the sites close to selected sites (Phung et al. 2016). At the chromosome level, at least in humans, both gene conversion and linked selection may therefore contribute to the higher diversity reported in smaller chromosomes, though the generality of these findings is still limited to humans and few other species.

How can variation in mean recombination rate explain differences in divergence across chromosomes of varying size? The relationship between recombination and divergence could be explained by variation in ancestral polymorphism. Selection and/or mutagenic recombination in the ancestral population may lead to variation in genetic diversity across the genome (see above), and when this population splits into two populations, the initial differences between these daughter populations simply reflect patterns of diversity present in the parent population. Heterogeneous levels of diversity across the genome of ancestral populations hence may give rise to variation in divergence among chromosomes of different sizes between daughter populations. Alternatively, higher recombination could increase divergence by leading to the accumulation of more mutations between diverging populations, and hence affect divergence directly. As mutagenic and selective forces before and after splitting are not mutually exclusive and could both explain the effect of recombination on divergence (Kulathinal et al. 2008), it is crucial to understand the factors that shape the distribution of diversity across the genome.

The analysis of chromosome-level patterns of diversity and divergence are now possible thanks to the increasing number of high-quality, chromosome-level assemblies available for several closely-related species within a clade. Great apes and *Mus* mice were the first mammalian clades with sufficient genomic resources to enable genome-scale comparative genomics analyses (Thybert et al. 2018). For example, while within humans and among great apes, fine-scale diversity and divergence seem to be correlated with recombination, and recombination with chromosome size (Hellmann et al. 2003; Jensen-Seaman et al. 2004), these relationships are weaker or not present in the house mouse *Mus musculus* (Jensen-Seaman et al. 2004; Kartje et al. 2020). Great apes and *Mus* mice are phylogenetically distant, with an estimated last common ancestor ~90 million years ago (timetree.org), and differ in genome size, 3.1 Gb and 2.6 Gb for the human and the *Mus musculus* genomes, respectively. Furthermore, they differ substantially in their degree of genome structure conservation, with only one major chromosomal fusion in humans compared to the other great apes (2n=46-48) and extreme variation in chromosome number in *Mus* mice (2n=22-48; Hauffe and Searle 1993). Although rodents of the genus *Peromyscus* look similar to *Mus* mice in appearance and have similar genome sizes, they last shared a common ancestor with *Mus* ~25 million years ago (Steppan et al. 2004) and have very karyotypically stable genomes (2n=48; Smalec et al. 2019), even more so than great apes. As a rodent lineage with increasing genomic resources (Colella et al. 2020; Tigano et al. 2020; Colella et al. 2021) and conserved genome structure, *Peromyscus* offers similarities and contrasts to both the *Mus* and the great apes lineages, thus representing an ideal third clade to understanding the role of genome structure stability and the relationships between chromosome size, recombination, diversity, and divergence in mammals. For example, similar to the great apes (Hellman et al. 2003) but contrasting with *M. musculus*, the cactus mouse (*Peromyscus eremicus*) shows a strong inverse correlation between chromosome size and *π* (Tigano et al. 2020); conserved synteny, recombination rates, and crossover patterning among *Peromyscus* species (Peterson et al. 2019; Smalec et al. 2019) together suggest that this relationship between chromosome size and *π* may be common among species in this genus. In light of these observations on the correlation (or lack thereof) between *π* and chromosome size, we hypothesize that also *d* will show a negative relationship with chromosome size in great apes and *Peromyscus* but not in *Mus*.

By combining the analysis of chromosome-level genome assemblies from three different mammalian clades -*Mus* spp., *Peromyscus* spp., and great apes - and individual-based evolutionary simulations, we tested a) whether chromosomes of different sizes show different levels of sequence divergence among species within a clade and b) the evolutionary, demographic, and molecular factors linking recombination, diversity within species, and divergence among species. Through evolutionary simulations, we tested the role of recombination, effective population size *N_e_*, severity of bottleneck associated with population splitting, gene conversion, selection, divergence time, and the interplay of these factors in generating and maintaining genetic diversity and divergence. Our results show that heterogeneous recombination rates across the genome and their interplay with selection and demographic factors affect genetic diversity and divergence not only at a small scale within a chromosome, but also at a broad scale among chromosomes of different lengths, thus providing new and important insights into the role of genome structure, and potentially chromosomal rearrangements, in the heterogeneous distribution of genetic diversity and divergence within and between species. These findings are foundational to our understanding of the process of speciation and adaptation, and highlight the importance of considering the structure and the heterogenous distribution of recombination rates of the genome for the inference of selective sweeps and demographic histories.

## Methods

### Analyses of divergence

We examined chromosome-level reference genomes for four *Mus* species (*M. musculus*, *M. spretus*, *M. caroli*, and *M. pahari*), five great apes (*Homo sapiens, Pan troglodytes, Pan paniscus, Gorilla gorilla*, and *Pongo abelii*), and six *Peromyscus* species (*P. maniculatus*, *P. polionotus*, *P. eremicus*, *P. crinitus*, *P. nasutus*, and *P. californicus*; Accession numbers in Table S1). Reference genomes for all these species were publicly available, except for *P. nasutus* and *P. californicus*, which we *de novo* assembled using a combination of sequencing approaches and final chromosome-scaffolding with Hi-C data. For *P. nasutus*, we obtained a tissue sample from a female individual collected at El Malpais National Conservation Area (New Mexico, USA) and stored at the Museum of Southwestern Biology (MSB:Mamm:299083). We extracted high molecular weight DNA with the MagAttract HMW DNA Kit (QIAGEN) and selected long fragments (> 10 kb and progressively above 25 kb) using a Short Read Eliminator Kit (Circulomics Inc.). A 10X Genomics linked-read library was generated using this high-quality DNA sample at Dartmouth Hitchcock Medical Center (New Hampshire, USA) and sequenced at Novogene (California, USA) using one lane of 150 bp paired-end reads on an Illumina HiSeq X sequencing platform. We produced a first draft of the *P. nasutus* genome assembly using *Supernova 2.1.1* (Weisenfeld et al. 2017) with default settings and these linked reads as input. To order and orient scaffolds in chromosomes we generated and sequenced a proximity-ligation library (Hi-C) from the same sample used for the 10X library as part of the DNA Zoo consortium effort (dnazoo.org). The Hi-C data were mapped to the 10X assembly with Juicer (Durand et al. 2016) and scaffolds were ordered and oriented in chromosomes with the *3D-DNA* pipeline (Dudchenko et al. 2017) and Juicebox Assembly Tools (Dudchenko et al.). The Hi-C data are available on www.dnazoo.org/assemblies/Peromyscus_nasutus and can be visualized using Juicebox.js, a cloud-based visualization system for Hi-C data (Robinson et al. 2018). For the *P. californicus* genome, high molecular weight DNA was extracted from liver tissue from a captive female individual from a colony maintained at the University of Maryland (USA) and sequenced using 10X Genomics technology at the UC Davis Genome Center (California, USA). A first draft genome for *P. californicus* was based on these 10X linked reads and assembled using *Supernova* as for *P. nasutus*. Then, Chicago and Dovetail Hi-C libraries were created by Dovetail Genomics (California, USA) and used to scaffold the draft assembly with the HiRise pipeline. The Chicago data were used first and the resulting improved assembly was used as input for a second round of scaffolding with the Hi-C data only. The alignment of several *Peromyscus* genomes revealed some assembly errors in the existing *P. eremicus* assembly (Tigano et al. 2020), so we generated an additional Hi-C library from a primary fibroblast collection at the T.C. Hsu Cryo-Zoo at the University of Texas MD Anderson Cancer Center. Using the new data, we performed misjoin correction and re-scaffolding using 3D-DNA (Dudchenko et al., 2017) and Juicebox Assembly Tools (Dudchenko et al., 2018). The new Hi-C data for *P. eremicus* are available on www.dnazoo.org/assemblies/Peromyscus_eremicus and are visualized using Juicebox.js (Robinson et al. 2018). Although the misassemblies present in the previous *P. eremicus* genome assembly were intrachromosomal and did not greatly affect estimates of sequence divergence at the chromosome-level, here we report divergence estimates based on the more accurate assembly.

We generated pairwise alignments and estimated sequence divergence (*d*) with *Mummer4* (Marçais et al. 2018) and custom scripts (https://github.com/atigano/mammal_chromosome_size). First, we aligned pairs of genomes in each clade with *nucmer*, randomly choosing one as the reference and the other as the query, and the settings *--maxgap 2000* and *--mincluster 1000*. We retained a global set of alignments (*-g*) longer than 10 kb using *delta-filter* and converted the output into ‘btab’ format using *show-coords*. To identify and exclude N-to-N matches from downstream analyses we based our analyses on the estimated ‘percent similarity’ rather than ‘percent identity’. As percent similarities were calculated for alignments of different lengths, we calculated weighted mean chromosome-level *d* (= 1 - (percent similarities)) for each chromosome correcting for alignment length. For the purpose of this study, we focused on autosomes and excluded estimates for sex chromosomes, when present in the genome assembly, because sex chromosomes experience a different combination of evolutionary forces than do autosomes. We tested the ability of the log10-transformed chromosome size in bp (explanatory variable) to predict mean chromosome-level divergence (response variable) separately for each species pairwise alignment using linear models (simple linear regressions), and plotted these relationships in R version 3.6.2 (R core team).

### Evolutionary simulations

To disentangle the factors contributing to the relationship between chromosome size, recombination, diversity and divergence, we performed individual-based time-forward evolutionary simulations in SLiM3 (Haller and Messer 2019). We simulated individuals using a Wright-Fisher model, where an ancestral population (popA) splits into two populations (pop_1_ and pop_2_) after 20*N_e_* generations (Fig. S1A), and prevented gene flow between diverging populations pop_1_ and pop_2_ to control for this potential confounding factor. We selected this time as a burn-in to allow for coalescence, to generate diversity and to reach stable allele frequencies. To test for the effect of population size and its changes over time, we simulated ancestral populations pop_A_ of 10,000, 40,000 and 160,000 individuals, which grossly encompass variation in *N_e_* among great apes, *Peromyscus* and *Mus* (Lack et al. 2010; Phifer-Rixey et al. 2012; Prado-Martinez et al. 2013; Harris et al. 2016; Colella et al. 2021) and modeled bottlenecks of different severity associated with the split of pop_A_ into two daughter populations pop_1_ and pop_2_: individuals from pop_A_ were either sorted into two daughter populations of equal size (*N_e_* in pop_1_ and pop_2_ were 0.5*N_e_* of pop_A_) or an additional bottleneck further reduced *N_e_* in pop_1_ and pop_2_ to 0.1*N_e_* of pop_A_. As our working hypothesis was that chromosome size affects diversity and divergence due to higher recombination rates *r* in smaller chromosomes, we simulated chromosomes of fixed length (1 Mb) with varying *r* to account for chromosome size variation, while keeping everything else the same. Assuming one crossover/chromosome on average (Peterson et al. 2019), mean chromosome-wide *r* was 10^−8^, so to encompass variation in chromosome size in the mammals examined we simulated nine different recombination rates, spanning 0.33*r* to 3*r*, which extends beyond the variation in recombination rates expected to occur in these species based on variation in chromosome size. Note that recombination rates are even across the chromosome and constant through time. We calculated mean gene size (including introns and exons) and mean distance between genes from the gene annotation of the *P. eremicus* genome and built the chromosome structure based on these values, resulting in each chromosome having 9 coding genes of 20.5 kb separated by 94.5 kb of intergenic sequence (Fig. S1B). We used a uniform germline mutation rate as estimated in *M. musculus* (5.7*10^−9^; Milholland et al. 2017) across all simulated chromosomes and models. We modeled gene conversion rate (*r*/3) and gene conversion tract length (440 bp), when included in the model, based on estimates in *Drosophila melanogaster* (Miller et al. 2016), as no mammal-specific estimates have been established. In neutral models all mutations were neutral, whereas in models with selection mutations in coding genes could be neutral, deleterious or advantageous at a relative frequency of 0.3/1/0.0005, with the non-neutral mutations being always codominant. The fitness effects of the non-neutral mutations were drawn from a gamma distribution with a mean selection coefficient *s* of ±15.625*10^−3^ and a shape parameter alpha of 0.3 based on the parameter space explored by Campos and Charlesworth (2019) and Stankowski et al. (2019). We scaled *N_e_*, *μ*, and *r* by a factor of 25 to expedite simulations and ran 30 unique simulation replicates for each combination of parameters. All the different parameters used in the simulations are summarized in Table 1. Finally, to validate our assumption that varying recombination rate in lieu of chromosome length recapitulates differences among chromosomes of different lengths, we also simulated three additional chromosomes whose length matched the respective recombination rates. To maintain the proportion of coding sequence constant among chromosomes of fixed and varying length, chromosomes that were longer or shorter than 1 Mb had proportionally more or less coding genes, respectively. We used the 1 Mb chromosome with 1*r* as reference and added a chromosome of 3 Mb with 0.33*r*, the lowest recombination rate simulated, a chromosome of 0.33 Mb with 3*r*, the highest recombination rate simulation, and a third chromosome of 0.66 Mb chromosome with 1.5*r*.

**Table 1.** Summary of parameters used in evolutionary simulations.

| Variable | Values | Scaled values as in simulations (x25) |
| --- | --- | --- |
| Mutation rate | $5.7 \times 10^{-9}$ | $1.42 \times 10^{-7}$ |
| Mean recombination rate $r$ | $10^{-8}$ | $2.5 \times 10^{-7}$ |
| Gene conversion rate | $r/3$ | $r/3$ |
| Gene conversion tract length | 440 bp | 440 bp |
| Selection coefficient $s$ | $\pm 15.625 \times 10^{-3}$ | $\pm 15.625 \times 10^{-3}$ |
| Relative frequency of neutral, deleterious, and advantageous mutations | 0.3, 1, and 0.0005 | 0.3, 1, and 0.0005 |
| Selection model | neutral/with selection | neutral/with selection |
| $N_e$ of pop<sub>A</sub> | 10,000/40,000/160,000 | 400/1,600/6,400 |
| Reduction of $N_e$ of pop<sub>1</sub> and pop<sub>2</sub> relative to pop<sub>A</sub> | 0.5/0.1 | 0.5/0.1 |
| Recombination rates | $0.33r/0.4r/0.5r/0.66r/r/1.5r/2r/2.5r/3r$ | $0.33r/0.4r/0.5r/0.66r/r/1.5r/2r/2.5r/3r$ |
| Gene conversion | yes/no | yes/no |

To investigate the factors affecting levels of diversity in chromosomes of different sizes, we sampled 30 individuals when pop_A_ reached 20*N_e_* generations (i.e. right before the split) for each simulation, output variant sites in a VCF file, and calculated *π* across the chromosome, in coding genes only, and in intergenic areas only, using VCFtools (Danecek et al. 2011). To obtain estimates of sequence divergence from simulations comparable to those from pairwise alignment of genome assemblies, we calculated *d* as the proportion of unmatched bases between two haploid genomes sampled randomly from each of the two diverging populations pop_1_ and pop_2_. We output these estimates right after the split and every 250,000 generations afterwards, up to 10 million generations. To further disentangle the effect of direct and linked selection, we estimated *d* also in coding genes and intergenic areas separately between the same genomes sampled above one generation after the split, when *d* is highest and not affected by decay yet (see Results and discussion). All scripts used in simulations are available at https://github.com/atigano/mammal_chromosome_size/simulations/.

## Results and discussion

### Empirical data show a strong, inverse relationship between chromosome size and divergence between species, with a few exceptions

Chromosome size was a strong significant predictor of mean sequence divergence *d* in each of the species pairwise comparisons in *Peromyscus* and great apes (all comparisons had *p* < 0.001 using linear models; Fig. 1A and 1C for example comparisons). Chromosome size showed a negative relationship to mean *d* and explained 62-89% and 46-65% of the variance in mean *d* across chromosomes (all *R*^2^ are adjusted hereafter) in *Peromyscus* and great apes, respectively. Among *Mus* spp., we found a significant, negative relationship between chromosome size and *d* only between *M. pahari* and *M. spretus* (*p* < 0.001; Fig. 1E), which explained 42% of the variance in *d* across chromosomes. We hypothesized that the discrepancy between results from *Mu*s and the other two clades examined could be explained by the relatively poor genome structure conservation in *Mus*, so we investigated this further. Among the *Mus* genome alignments, the *M. pahari*/*M. spretus* comparison was the only one where *M. pahari* was used as a reference genome. *M. pahari* is the most divergent (3-6 million years ago) and differs from the other *Mus* species in that it shows a karyotype with 24 chromosomes, while *M. spretus, M. musculus* and *M. caroli* exhibit karyotype with only 20 chromosomes (Thybert et al. 2018). Further, fewer synteny breaks between *M. pahari* and the rat (*Rattus norvegicus*) relative the other *Mus* species analyzed here (19 versus 35) demonstrate that the *M. pahari* karyotype is the most similar to the ancestral karyotype of the *Mus* species included here (Thybert et al. 2018). As the reference and query genomes in each pairwise comparison were chosen randomly, we produced new alignments and calculated *d* for all the possible pairwise combinations within each clade to test for the effect of the reference genome to chromosome-level *d* estimates (Fig. S2, S3, S4). While in the *Peromyscus* and great apes clades all reference-query combinations, including the reciprocal of the comparisons first analyzed (Fig. 1B and 1D) showed a strong negative relationship between chromosome size and *d* (R^2^= 0.59-0.91 among *Peromyscus* and 0.46-0.65 among great apes, all *p* < 0.001; Fig. S2 and S3), in the *Mus* clade this relationship emerged only when *M. pahari* was used as reference genome (p < 0.001; Fig. 1F and S4) and explained 52 and 54% of the variance in *M. musculus* and *M. caroli*, respectively (Fig. S4). While a significant positive correlation between neutral human-primate divergence and human recombination rate has been reported at smaller scales (i.e. in 100 kb sliding windows across the genome; Phung et al. 2016), our results show that these relationships between recombination, diversity, and divergence are strong at a large, chromosome scale within three different highly divergent clades (~30-90 MYA), including *Mus*, where, at least in *M. musculus*, the relationship between π and recombination at smaller scales is not always significant (Kartje et al. 2020).

**Fig.1.**
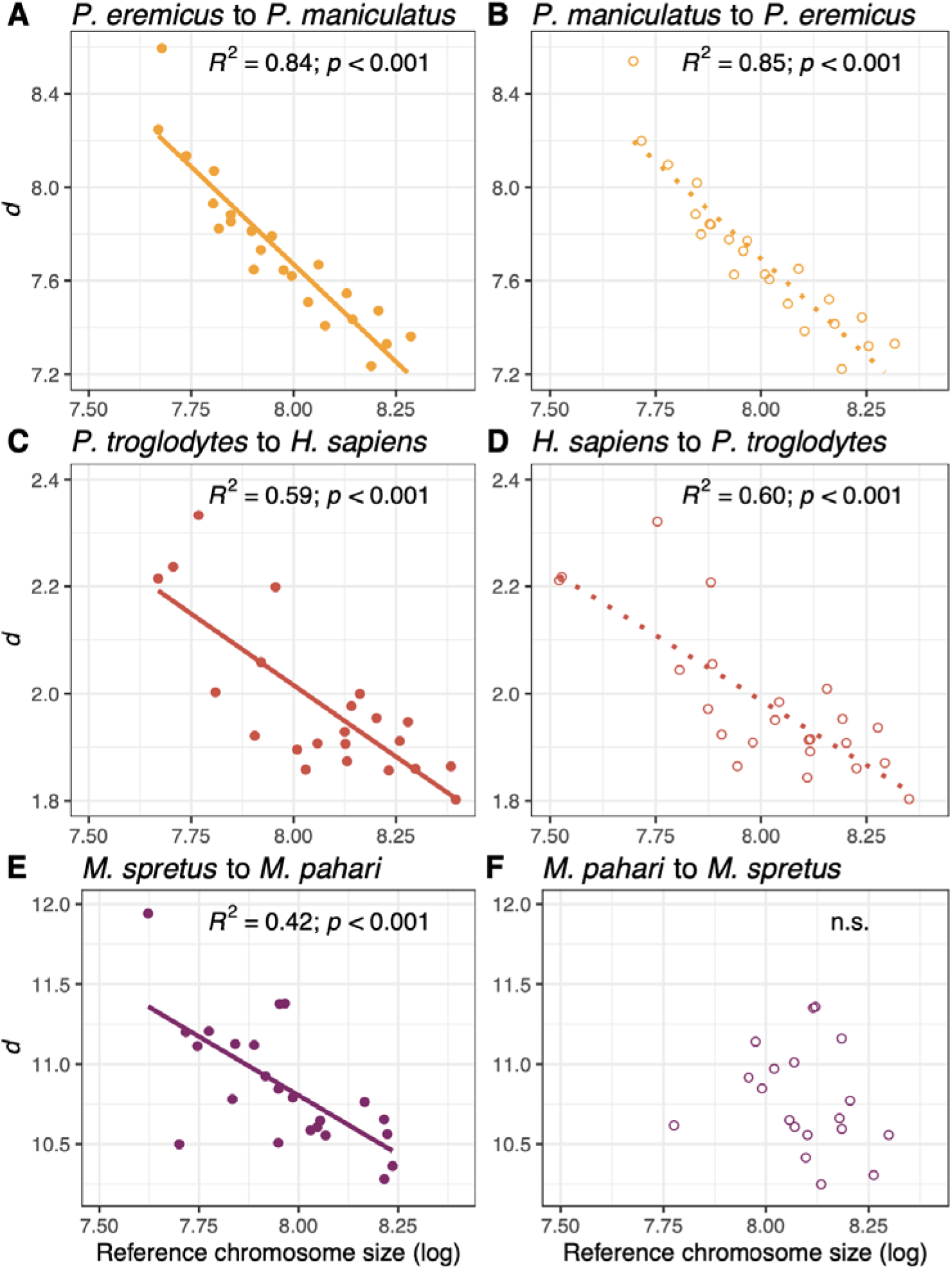
Plots showing the relationship between log10-transformed chromosome size (bp) and sequence divergence among species within the *Peromyscus*, Hominidae and *Mus* clades. On the left panel (A, C, E), one representative comparison from each of the *Peromyscus*, Hominidae, and *Mus* clades are displayed (see Figure S2, S3, and S4 for all comparisons). The comparison of the same species pairs are represented on the right panel but the query and reference species are inverted in plots B, D, F to highlight that in the *Mus*, but not in *Peromyscus* and *Hominidae* clades, the choice of the reference genome affects the correlation between chromosome size and *d*. In the bottom panel, the comparison between *Mus spretus* and *M. pahari* is shown, with *M. pahari* as reference on the left (E) and with *M. spretus* as reference on the right (F).

The fact that chromosome size in the most ancestral karyotype (*M. pahari*), but not the more derived karyotypes (*M. musculus*, *M. spretus*, and *M. caroli*), is a strong predictor of levels of *d* between species with different genome structures indicates that these patterns evolve and are maintained across long evolutionary scales. The retention of ancestral patterns suggest that recombination hotspots could be conserved in rearranged chromosomes despite the evolution of a different genome structure, or that a different genomic landscape of recombination has not been sufficient to redistribute variation in sequence divergence expected based on differences in chromosome size over this time scale. In *Heliconius* butterflies, for example, patterns of diversity in fused chromosomes seem to have aligned to expectations based on the size of the derived chromosome size over time, rather than maintaining patterns consistent with the unfused chromosomes of origin (Cicconardi et al. 2021). Though the human genome underwent a chromosomal fusion compared to the other great apes, the correlation between chromosome size and *d* among great apes did not seem affected by the use of the human genome as reference (i.e. chromosome size did not explain a lower proportion of the variance in these comparisons; Fig. 1C, 1D, S3).

The choice of a model species is largely based on its potential to provide insights that are generalizable to other organisms, yet our results builds on previous work in *M. musculus* (Jensen-Seaman et al. 2004; Kartje et al. 2020) showing that the *Mus* clade is rather an outlier, and does not serve as a good model to analyze general patterns with regards to how genome and chromosome structure affect and shape heterogenous levels and patterns of diversity and divergence across the genome. However, our analysis of the *Mus* genomes provides insight into the reasons a species may deviate from expectations, showing that the dramatic chromosomal rearrangements occurred in the *Mus* clade seem to explain the apparent lack of a relationship between chromosome size and recombination rate, π, and *d* when a derived karyotype is used as reference. Moreover, the lower proportion of the variation in *d* explained in *Mus* compared to the other two clades, even when the ancestral karyotype of *M. pahari* is used as reference, may be due to an ongoing shift towards expectations based on the size of the rearranged chromosomes, as suggested in *Heliconius* butterflies (Cicconardi et al. 2021).

Another factor that might explain variation among clades in the strength of the association between chromosome size and divergence (Fig. 1) is gene flow and introgression. Introgression can increase diversity within species and decrease divergence between hybridizing species (Tigano and Friesen 2016); and the distribution and length of introgressed sequence from one species to another is determined by the interplay of selection and recombination (Duranton et al. 2018). It is therefore plausible that chromosomes of different sizes could affect the fate of introgressed DNA, in terms of both the distribution and length of donor sequence. Inversely, the heterogeneous landscape of introgression could affect the power of chromosome size in explaining variation in diversity and divergence across chromosomes. The adaptive introgression of a rodenticide-resistant allele from *Mus spretus* to *M. musculus* shows that the fitness benefit of a beneficial donor allele can overcome incomplete reproductive barriers between species (Song et al. 2011). Moreover, the reticulate evolution of primates, and possibly many other understudied clades, support the occurrence of ancient gene flow and introgression among many extant and extinct taxa during and post-speciation (Feder et al. 2012; Vanderpool et al. 2020). However, although gene flow between these and with other unsampled species cannot be excluded, hybridization is rare among *Peromyscus* species (e.g., Barko and Feldhamer 2002; Leo and Millien 2017); hybrid sterility or infertility in *Mus* has been reported between species (Dejager et al. 2009) and even between *M. musculus* subspecies (Turner et al. 2011); and among great apes recent hybridization appears limited to intraspecific gene flow (Fontsere et al. 2019). Therefore, gene flow does not seem to affect the relationship between chromosome size and divergence among the species included in this study, though is a factor to consider when comparing species connected by historical or contemporary gene flow.

### Evolutionary simulations reveal the factors driving the empirical patterns

Simulations helped generate a mechanistic understanding of most of the empirical patterns reported in this and other studies (Dutoit et al. 2017; Murray et al. 2017; Kartje et al. 2020; Tigano et al. 2020). Patterns of diversity observed across simulated chromosomes that varied in recombination rate only were qualitatively equivalent to the simulated chromosomes that varied also in length (Fig. S6), showing that, as long as all the other parameters are constant and uniform across the chromosomes, including mutation rate and proportion of coding genes, variation in recombination rate alone suffices to recapitulate variation in chromosome length. Therefore, here we show and discuss results based on simulated chromosomes that vary in recombination rate only. In neutral simulations, we did not observe variation in *π* among chromosomes with different recombination rates whether the model included gene conversion or not (ANOVA, p > 0.05). These results indicate that recombination alone does not explain variation in *π* among chromosomes and that gene conversion does not contribute substantially to increasing levels of *π*, at least at the rates that we assumed and over relatively short evolutionary times (20*N_e_* generations). Gene conversion occurs at a fraction of the recombination rate and affects only a small segment of DNA (100-2000 bp) at a time (Korunes and Noor 2017; Korunes and Noor 2019), hence its effect on chromosome-wide levels of *π* may be detectable only over long evolutionary times. Recombination could also be mutagenic *per se* by promoting *de novo* mutations at the DNA breaks caused by crossovers, but the mechanism underlying this phenomenon is not clear (Hodgkinson and Eyre-Walker 2011). We did not model crossover mutagenesis in our simulations, but a recent study in humans found the mutation rate associated with crossovers to be ~4%, i.e. one *de novo* mutation every ~23 crossovers (Halldorsson et al. 2019) suggesting that crossover mutagenesis could contribute to, but not entirely account for, the variation in π among chromosomes of different sizes over long evolutionary times, similar to gene conversion. In contrast, in models with selection, recombination rate was a significant (p < 0.001) predictor of differences in *π* among chromosomes across populations of three vastly different *N_e_* (Fig. 2A). However, Δ*π* - the difference in *π* between the chromosomes with the highest and lowest recombination rates - spanned over two orders of magnitude when comparing the smallest and the largest simulated ancestral populations (Δ*π* = 3.16*10^−5^−2.69*10^−3^; Fig. 3). Also the proportions of variance in *π* explained by recombination rate increased with *N_e_*: they were 13, 60, and 85% in populations of 10,000, 40,000 and 160,000 individuals respectively in models without gene conversion (estimates were similar for models with gene conversion, except for the smallest *N_e_* where *r* explained 27% of the variation), suggesting that while selection is the main determinant of the relationship between recombination and π in large populations, genetic drift prevails in smaller populations.

**Fig. 2.**
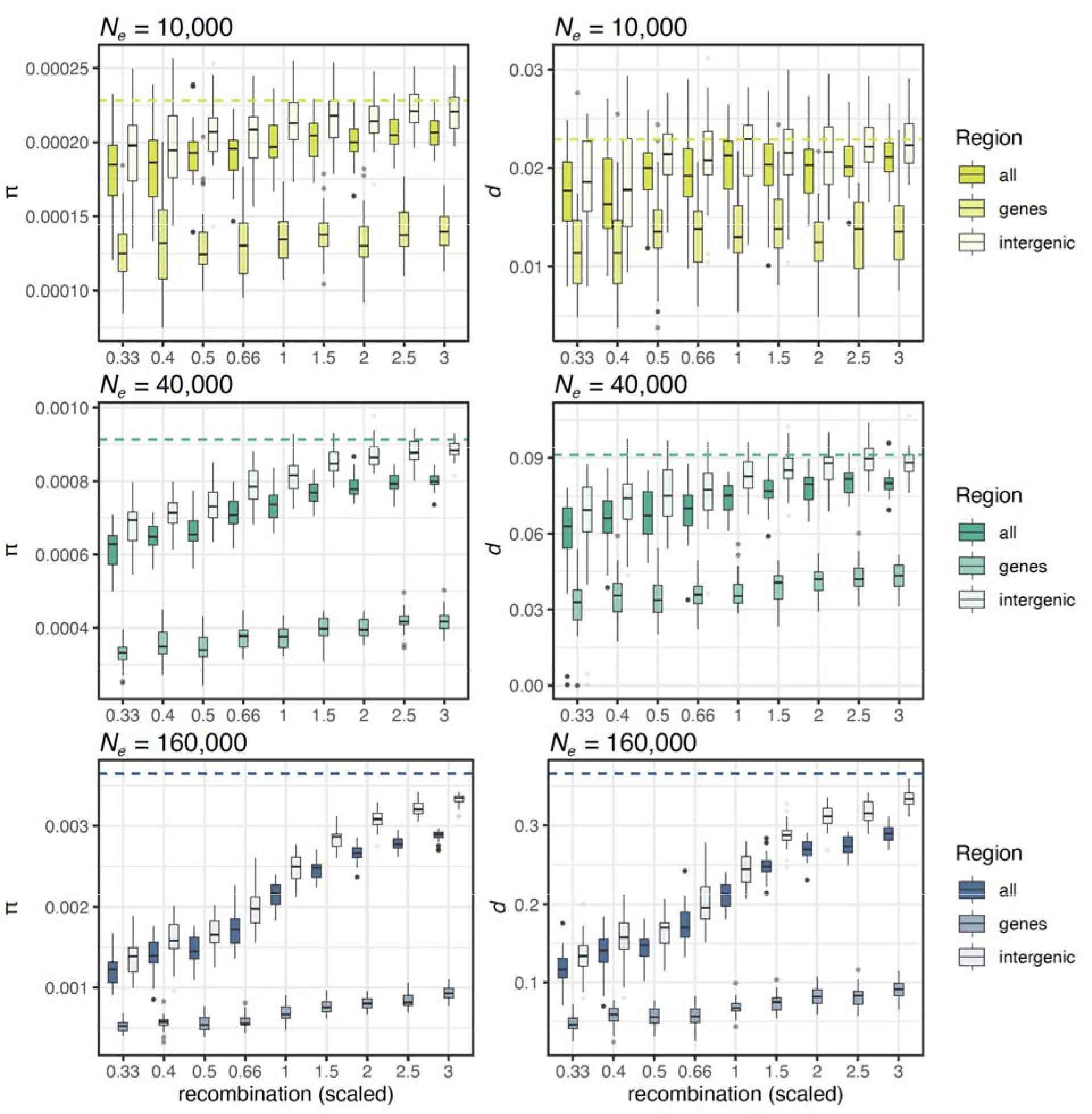
Boxplots summarizing results from evolutionary simulations on the relationship between recombination rate and *π* after 20*N_e_* generations in the ancestral population (panel A on the left) and *d* one generation after the split after a mild bottleneck (0.5 of ancestral *N_e_*) between the two daughter populations (panel B on the right) in each of three simulated ancestral *N_e_*. Boxplots refer to the results from the models with selection and the dashed line shows the results from the neutral models. Here are the results with models without gene conversion as no significant differences were found between models with and without gene conversion.

**Fig. 3.**
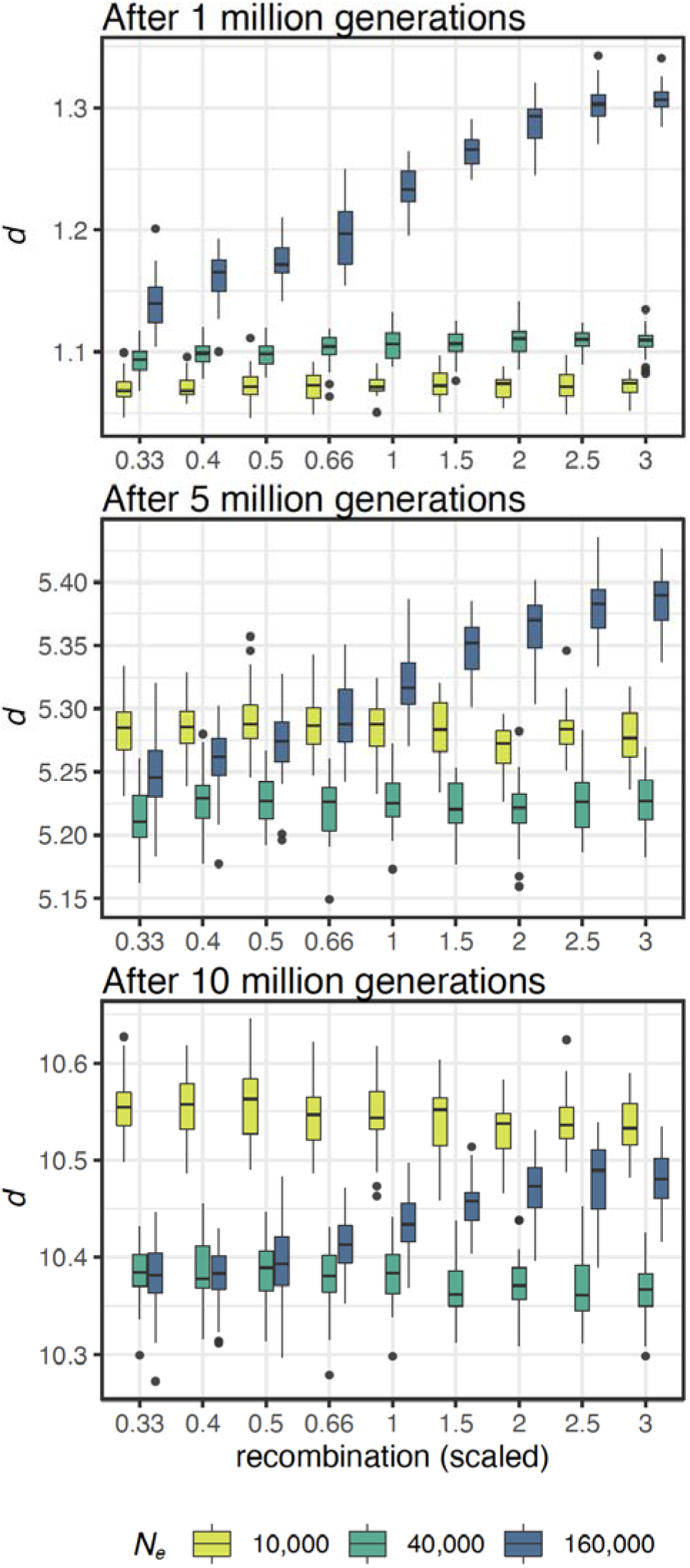
Boxplots summarizing results from evolutionary simulations on the relationship between recombination rate and *d* in models with selection and without gene conversion in each of three simulated ancestral *N_e_* and three time points after the split from the ancestral popA and a mild bottleneck. Gene conversion was not included in these models as no significant differences were found between models with and without gene conversion. (See Fig. S5 for comparisons with neutral models and models with a more severe bottleneck).

The comparison of *π* across models with and without selection shows that diversity is lower overall, regardless of the recombination rate, in chromosomes affected by selection (Fig. 2A). The reduction in *π* is strongest at the coding genes, which experience both positive and negative selection directly, and indirectly through linked selection (Fig. 2A). Chromosome-wide estimates are more similar to those based on the analysis of intergenic areas subjected to linked selection only (Fig. 2A), which is at least partly due to the relatively much larger proportion of non-coding over coding regions. Nonetheless, a positive relationship between recombination rate and π is evident across the different genomic areas and *N_e_* considered (Fig. 2A). The comparison of *π* estimates from across the chromosome, coding genes only, and intergenic areas only, corroborates that differences in diversity among chromosomes of different sizes are due to the balance between selection and recombination, with selection reducing diversity and recombination reducing linkage disequilibrium, which in turn reduces the effect of linked selection. As recombination increases, it more strongly counteracts linked selection in the intergenic areas, to the point of almost restoring levels of diversity like those expected under a neutral model (Fig. 2A). In other words, in our simulations the interplay between recombination and selection is the main factor driving the inverse correlation between chromosome size and *π*, a pattern that is described in many species.

The fact that recombination was a stronger contributor to variation in *π* in larger populations than in smaller ones could be due to one or the combination of two factors: 1) as the effect of selection in the genome depends on both effective population size and the strength of selection (*N_e_s*), larger populations will have proportionally more selective sweeps than smaller populations - because more mutations in small populations will have scaled coefficients so small that they will actually behave as neutral - and these sweeps will be more strong and efficient at removing diversity, and 2) larger populations will have higher population recombination rates (*ρ* = 4*N_e_r*), which will break linkage disequilibrium even more efficiently in chromosomes with high recombination rates. As smaller population will have proportionally more mutations behaving as neutral ones, more mutations will be more predominantly governed by stochasticity rather than by the deterministic effect of selection in these small populations (Charlesworth 2009), which is consistent with selection explaining less variation in *π* in smaller populations than in larger ones (see above).

In neutral models, *d* did not vary across chromosomes with different recombination rates (ANOVA, p >> 0.05; Fig. 3B), while in models with selection differences in *d* among chromosomes one generation after the split were significantly different from zero (ANOVA, p < 0.001; Fig. 2B) and strongly correlated with recombination rate (p < 0.001; Fig. 2B), i.e. chromosomes with lower recombination rates had lower *d* between populations, across all three simulated values of *N_e_*. Sequence divergence right after the split reflected the levels of diversity within the ancestral population before the split across all models, with higher Δ*d* - the difference in *d* between the chromosomes with the highest and lowest recombination rates - in larger populations (Fig. 2B). Similar to the patterns observed for π, *d* was lowest in coding regions and highest in intergenic areas, with chromosome-wide estimates lower than, but similar to, the latter (Fig. 2B). Testing empirically whether species with higher ancestral *N_e_* show higher Δ*d* would support the contribution of demography in the relationship between recombination and *d*. However, across the three clades examined here, *N_e_* and divergence times between species seem to covary, for example with great apes having not only the smallest *N_e_* but also the most recent species divergence, so that the actual relative contributions of *N_e_* and divergence times cannot be disentangled using empirical data in this study.

In neutral models, *d* increases linearly with time (4*N_e_μ* + 2T*μ*, where T is the number of generations) so the severity of the bottleneck at the time of the split does not have any effect on divergence between isolated populations in our neutral simulations, even at the smallest *N_e_*. Although genetic drift was *de facto* the only evolutionary force driving changes in allele frequency in these neutral models, its effect size (= 1/2*N_e_*) in our simulated populations was nonetheless very small (6.25*10^−6^ - 10^−4^), which indicates that new mutations are the main source of *d* over time in neutral models. In fact, our *d* estimates encompass both fixed and segregating mutations in each of the two compared populations.

In models with selection, *d* increased over time, though at a slower pace than in neutral models, and faster in smaller populations relative to larger populations (Fig. 3). Further, while larger populations showed higher overall divergence than smaller ones in the early stages of divergence, the trend reversed with time: after 10 million generations *d* between the smallest populations (*N_e_* = 5,000 individuals each) surpassed *d* both between the medium-sized (*N_e_* = 20,000 individuals each) and between the largest populations (*N_e_* = 80,000 individuals each; Fig. 3). This pattern was even more pronounced when populations pop_1_ and pop_2_ were affected by a stronger bottleneck at the time of the split from pop_A_: populations of all sizes accumulated *d* faster than in the models with a weaker bottleneck, and even faster between the smallest populations (*N_e_* = 1,000 individuals each) compared to estimates from larger populations (*N_e_* = 4,000 and 16,000 individuals each, respectively; Fig. S5). These results show how genetic drift is much stronger in small populations, but only in the models with selection (Fig. S5). Neutral mutations fix at a much faster rate than those under selection because the fixation probability of a locus under selection depends also on the strength of selection acting on its linked sites (Hill-Robertson interference; Hill and Robertson 1966; Felsenstein 1974). Given the distribution of fitness effects (DFE) we simulated, the probability of a beneficial mutation to effectively act as a neutral mutation (i.e. that *N_e_s* < 1) is higher in smaller populations than in larger ones (Charlesworth 2009). Therefore, in smaller population, a higher proportion of beneficial or deleterious mutations will act as neutral and their fixation probability will be higher and depend only on the combination of genetic drift and linked selection, which in turn will depend on the strength of selection acting on the linked mutation and the rate of recombination affecting linkage disequilibrium between the two mutations.

We found that Δ*d* decreased with divergence time in all models with selection (Fig. 4), suggesting that either divergence rate is relatively accelerated in large chromosomes or slowed down in small ones. Based on what discussed above, larger chromosomes, where recombination rates are lower, should experience stronger Hill-Robertson interference, and hence lower fixation probabilities and slower divergence rate, which is in contrast with what observed. Alternatively, lower recombination in larger chromosomes could strengthen the effect of linked selection, resulting in local chromosome-wide reductions in *N_e_*, and thus stronger genetic drift and faster rate of sequence divergence than smaller chromosomes with higher recombination rates. This is clearly illustrated by an empirical study on Bornean and Sumatran Orangutans (*P. pygmaeus* and *P. abelii*) showing that estimates of ancestral *N_e_*vary among chromosomes and that chromosome size is a strong predictor of variation in both inferred ancestral *N_e_* and recombination rate, which in turn suggests a direct relationship between these two (Mailund et al. 2011). Additionally, Phung et al. (2016) showed that the window-based correlation between recombination and divergence rate in simulated genomes decreased with splitting time, and attributed this decrease to a concurrent decrease in levels of ancestral variation. At the chromosome level, these observations suggest that chromosomes of different sizes may lose ancestral variation at different rates and thus may account for the decay in Δ*d*.

**Fig. 4.**
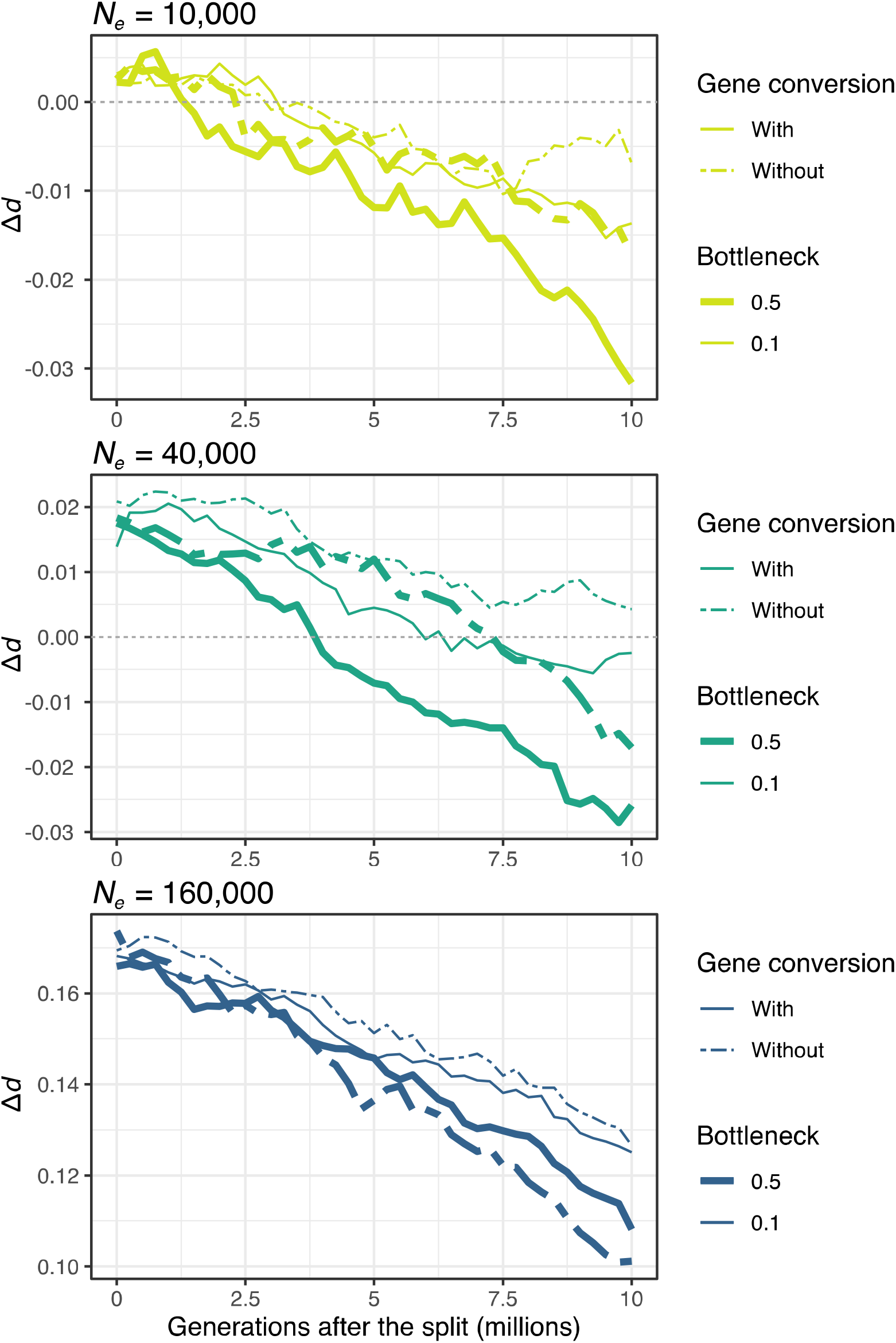
Plots showing the decay of Δ*d* over time in the evolutionary simulations based on the models with and without gene conversion, and with a mild (0.5) and a severe bottleneck (0.1) for each of the three simulated ancestral *N_e_*.

These different rates of divergence explain why over time Δ*d* becomes negative in our simulations, i.e. the chromosome with the lowest recombination rate becomes more divergent than the chromosome with the highest recombination rate, in the populations with the smaller *N_e_* (Fig. 4). Initially, the differences in Δ*d* will be determined by Δπ in the ancestral population at the time of the split, but with time the different divergence rates among chromosomes caused by the interplay of recombination, selection, and thus drift, will erode Δ*d*, and even reverse it (Fig. 4). Furthermore, the severity of the bottleneck at the species split affected the Δ*d* decay rate, with a faster decay in less severely reduced *N_e_*, regardless of ancestral *N_e_* (Fig. 4). Also gene conversion seemed to accelerate the Δ*d* decay overall, but not in the largest population experiencing the weaker bottleneck (Fig. 4). That Δ*d* decay was generally faster both in populations experiencing milder bottlenecks, which therefore had larger *N_e_*, and in models with gene conversion compared to those without, suggests that higher mutation rates in these cases can accelerate Δ*d* decay. Although the ultimate mechanism is not clear, faster Δ*d* decay may be explained by the rate of loss of ancestral polymorphism (Phung et al. 2016), which should be higher with higher mutation rates. Importantly, the effect of gene conversion becomes more evident after 5 million generations (Fig. 4), showing that, although gene conversion is not the main determinant of differences in π and *d* among chromosomes of different sizes, it can contribute to these patterns over long evolutionary times. The evolutionary simulations provide clear insights into why the variation in divergence among chromosomes of different sizes decreases with time, and suggest that it would require either small *N_e_* and/or long divergence times to observe a negative Δ*d*. We did not observe any negative Δ*d* in our species pairwise comparisons. In the future, the empirical test of these observations will require the inclusion of additional clades and larger number of species comparisons within clades to verify how commonly this occurs empirically and to disentangle the factors promoting, or hindering, this pattern.

Our simulations were based on reasonably realistic parameters, except for the absence of neutrally evolving introns in simulated genes, to reach an acceptable compromise between capturing the complexity of the evolutionary processes, and their interactions, while maintaining enough statistical power to understand the relative roles of the many factors at play. Notwithstanding that empirically-derived values for the factors included in our simulations are not available for most species or are difficult to accurately estimate, excluding introns provided more target sequence for selection to act on and a good trade-off in terms of computational time and resources. With the exception of a clear effect of *N_e_* and divergence time in the magnitude of the variation in *d* observed among chromosomes, our evolutionary simulations well illustrate the processes driving the empirical patterns of divergence between species described in three different clades of mammals: in the absence of chromosomal rearrangements, the interplay of recombination and selection determines levels of *π* in the ancestral species; higher *π* in the ancestral species results in higher *d* among haplotypes sorted into the daughter species, thus explaining the differences in *π* and *d* among chromosomes of different sizes.

### Empirical analyses and simulations highlight the rule and the exceptions

Species showing an inverse relationship between recombination rate and chromosome size are found among mammals, birds, yeast, worms, and plants (Pessia et al. 2012), which highlights that this relationship is not an idiosyncratic feature of a particular taxon but rather a widespread feature of genome evolution. Our evolutionary simulations show that varying recombination rates across chromosomes should result in differences in *π* and *d* among chromosomes of different sizes, but empirical support for this prediction is mixed. In *M. musculus*, for example, chromosome size is not a good predictor of variation in *π* within the species (Pessia et al. 2012) or *d* to other *Mus* spp. (our study). Our analyses here have shown that this lack of correlation is likely due to dramatic changes in the genome structure of *M. musculus* and other congeneric species relative to their common ancestor (most similar to *M. pahari*). Future work, including additional species comparisons and evolutionary simulations, will focus on testing the role of chromosomal rearrangement in the presence of a relationship between chromosome size and divergence across species and over time. These results also stress the importance of the choice of the reference genome in this type of analysis. Not only does the reference genome potentially mask existing relationships due to the evolution of different genome structure, as we have shown here, but also its quality is crucial to obtain high-quality genome alignments, to calculate *d* accurately, and to estimate chromosome sizes from sequence length in the absence of cytological data.

No significant relationship between chromosome size and divergence between human and chimpanzee was found in another study (Patterson et al. 2006), but these results were based on only 20 Mb of aligned sequences. Moreover, different avian species have shown a positive (Dutoit et al. 2017), a negative (Manthey et al. 2015), or no relationship (Callicrate et al. 2014) between chromosome size and π. Dramatic chromosomal rearrangements can be excluded in these examples (Ellegren 2010), begging the question: what other factors could explain these deviations from our model? First, given the variation in *d* within chromosomes, incomplete genome sampling may confound these chromosome-level relationships. Second, Dutoit and colleagues (2017) argue that in the collared flycatcher (*Ficedula albicollis*) a positive relationship between chromosome size and *π*, which is opposite to expectations, could be explained by the density of targets of selection, higher in smaller chromosomes than in larger ones in this species. However, given the high degree of synteny conservation among birds (Ellegren 2010), all avian species should show a similar pattern, which is not the case. For example, the comparison of genome-wide patterns in *π* in the passenger pigeon (*Ectopistes migratorius*), known as the most abundant bird in North America before it went extinct, and the band-tailed pigeon (*Patagioenas fasciata*), with a current population size three orders of magnitude smaller, not only shows higher *π* in smaller chromosomes in both species, but also the effect of *N_e_* on Δ*π*, as per our predictions (Murray et al. 2017). Our analyses highlight the contribution of demography (i.e. *N_e_*, severity of bottleneck and genetic drift) in affecting, and even reversing, the relationship between recombination, *π*, and *d*, which could potentially explain the opposite correlation reported in the collared flycatcher. The importance of historical demography has been demonstrated also in the divergence of the sex chromosome Z in *Heliconius* butterflies using a combination of empirical data and evolutionary simulations (Van Belleghem et al. 2018). Alternatively, the strength of selection, rather than the density of targets of selection, could disrupt the correlation between chromosome size and *π* in case of strong selective sweeps preferentially occurring in small chromosomes. Finally, limited variation in recombination, *π*, and *d* among chromosomes could be simply due to lack of variation in chromosome size. The analysis of 128 eukaryotic and prokaryotic genomes has shown that variation in chromosome size is directly proportional to genome size (Li et al. 2011), suggesting that variation in recombination, *π*, and *d* among chromosomes should decrease with genome size.

## Conclusions

Variation in recombination across the genome affects the evolution and maintenance of traits relevant to adaptation and speciation, the genomic architecture of the loci underlying those traits, and our ability to detect those loci (Yeaman and Otto 2011; Yeaman 2013; Cruickshank and Hahn 2014; Burri et al. 2015; Roesti 2018; Lotterhos 2019; Booker et al. 2020). Heterogenous recombination rates can also lead to the inference of different demographic histories and ancestral *N_e_* estimates depending on the chromosome or region of chromosome analyzed (Mailund et al. 2011; Robinson et al. 2021). We have shown strong evidence from empirical analyses and evolutionary simulations that the inverse relationship between recombination rate and chromosome size can result in significant differences in *π* and *d* among chromosomes of different sizes, indicating that variation in recombination rates among chromosomes of different sizes has an overall stronger effect than variation in recombination rates within chromosomes.

In the clades included in this study, *N_e_* covaries with divergence time scales, thus it is not possible to disentangle the relative effect of these two factors on patterns of divergence at this time. Furthermore, we cannot currently demonstrate that the chromosomal rearrangements in the *Mus* clade have a causal effect on masking the ancestral patterns of divergence, but nonetheless show a strong correlation. Future analyses of species and clades with different combinations of *N_e_*, divergence times, and degree of genome structure conservation will help address these gaps. Nonetheless, our study shows that chromosome size should be considered in the study of the genomic basis of adaptation and speciation. Do smaller chromosomes play a proportionally more prominent role than larger chromosomes in adaptation and speciation? Or are these differences in *π* and *d* strong enough to confound signals of selection in the genome? As chromosome-level assemblies and population whole genome resequencing data of closely-related species become available for an increasing number of taxa, the combination of empirical and theoretical investigations will help address these outstanding questions and generate new ones on chromosome and genome evolution.

