## Supplementary Material for "Chromosome size affects sequence divergence between species through the interplay of recombination and selection"

**Table S1.** Accession number or DNA zoo link to all genomes included in our analysis of sequence divergence

| **Species** | **NCBI Accession Number** | **DNAzoo link** |
| --- | --- | --- |
| *Peromyscus maniculatus* | GCA_003704035.3 |  |
| *Peromyscus polionotus* | GCA_003704135.2 |  |
| *Peromyscus eremicus* |  | https://www.dnazoo.org/assemblies/Peromyscus_eremicus |
| *Peromyscus californicus* | GCA_007827085.2 |  |
| *Peromyscus crinitus* |  | https://www.dnazoo.org/assemblies/Peromyscus_crinitus |
| *Peromyscus nasutus* |  | https://www.dnazoo.org/assemblies/Peromyscus_nasutus |
| *Pan troglodytes* | GCA_002880755.3 |  |
| *Pan paniscus* | GCA_000258655.2 |  |
| *Homo sapiens* | GCA_000001405.28 |  |
| *Gorilla gorilla* | GCA_008122165.1 |  |
| *Pongo abelii* | GCA_002880775.3 |  |
| *Mus musculus* | GCA_000001635.9 |  |
| *Mus spretus* | GCA_001624865.1 |  |
| *Mus caroli* | GCA_900094665.2 |  |
| *Mus pahari* | GCA_900095145.2 |  |

**Fig. S1.** Figure depicting the demographic model and the chromosome structure used in the evolutionary simulations.

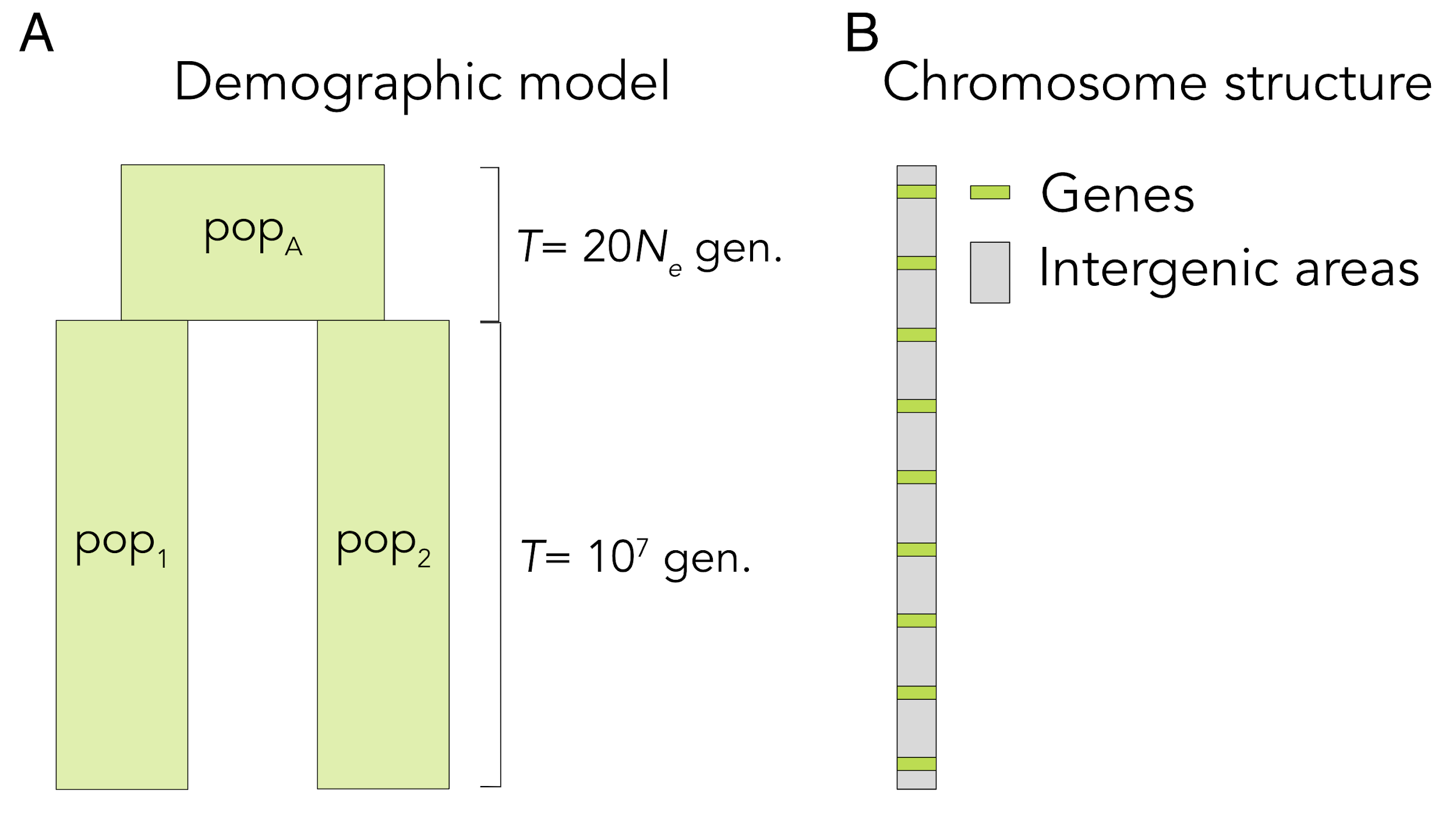

**Fig. S2.** Plots of the relationship between *d* and reference chromosome size (log-transformed) in *Peromyscus*. On the left are the reference genomes, on top the query genomes. All correlations are significant (*p* < 0.001).

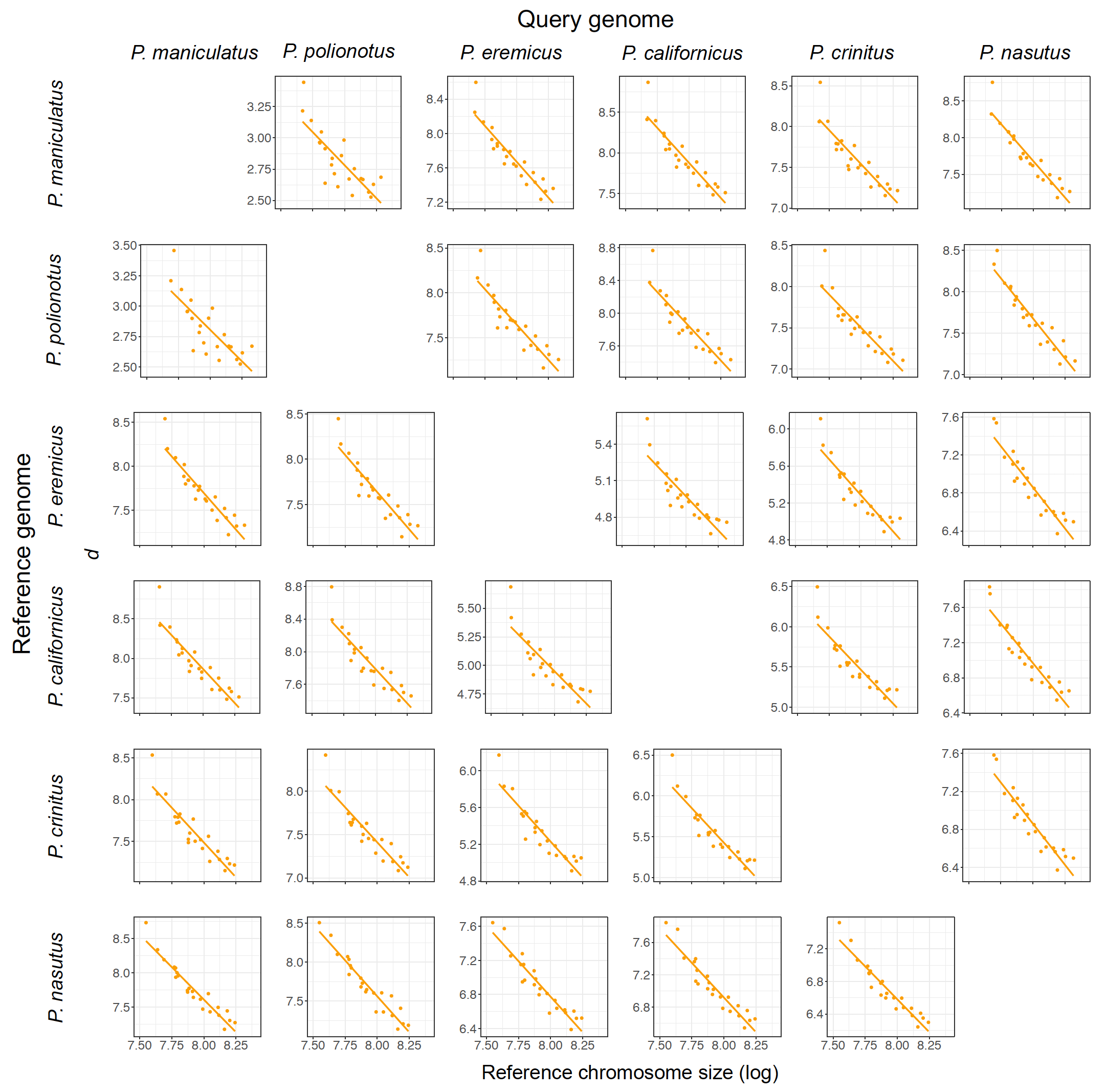

**Fig. S3.** Plots of the relationship between *d* and reference chromosome size (log-transformed) in great apes. On the left are the reference genomes, on top the query genomes. All correlations are significant (*p* < 0.001).

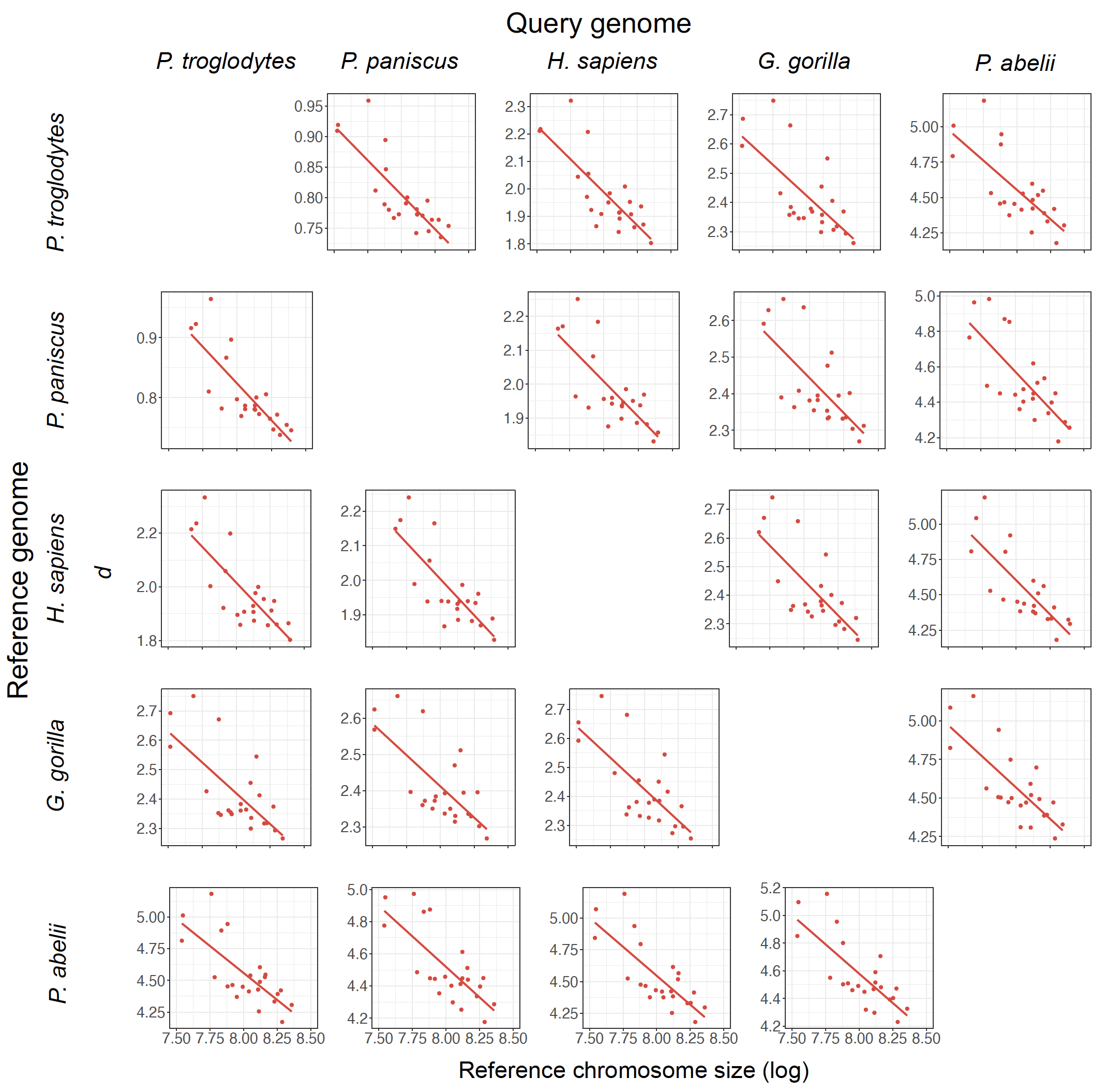

**Fig. S4.** Plots of the relationship between *d* and reference chromosome size (log-transformed) in *Mus*. On the left are the reference genomes, on top the query genomes. Correlations are significant (*p* < 0.001) only when genomes are aligned to *M. pahari* (bottom line).

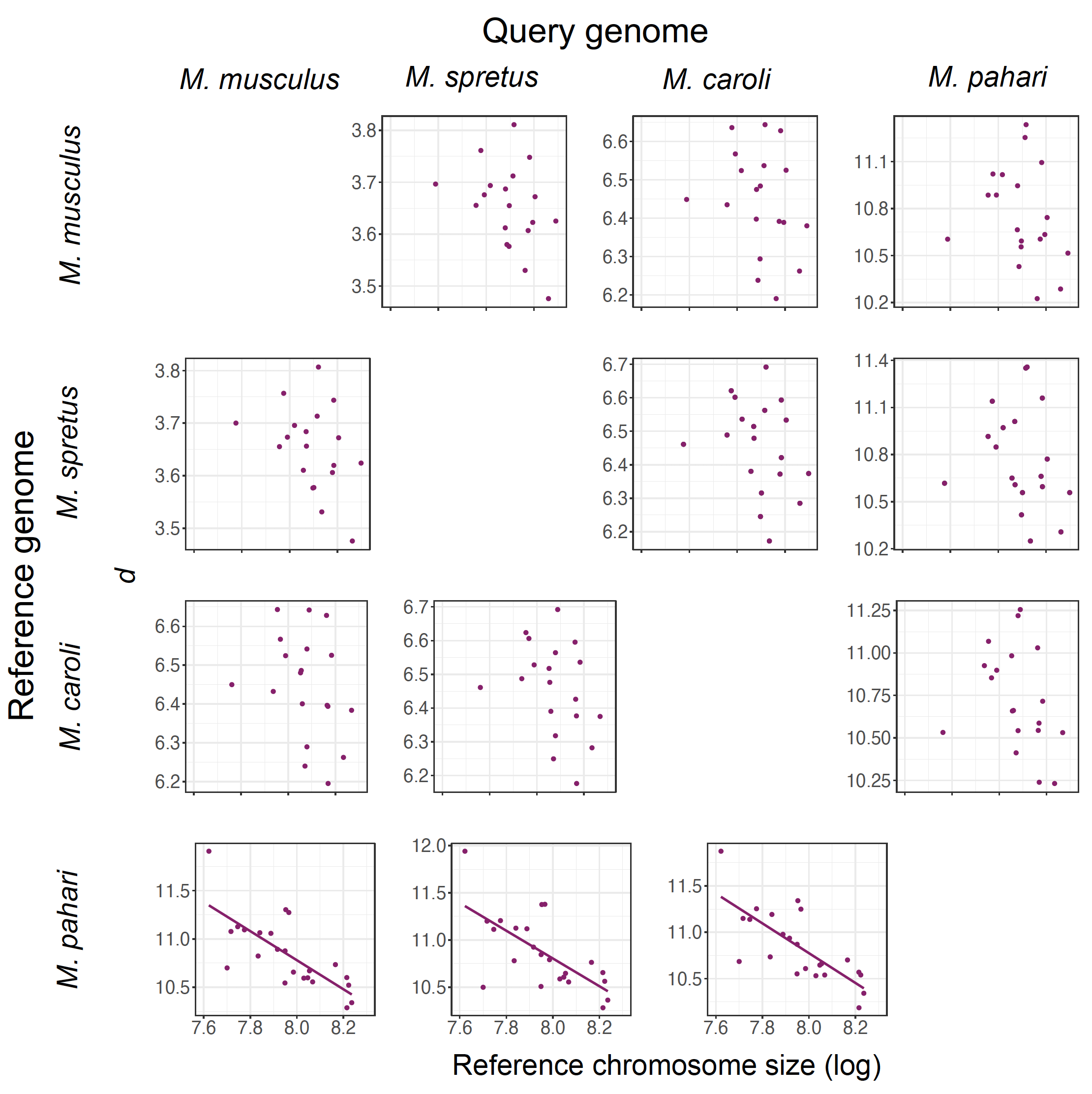

**Fig. S5.** Boxplots summarizing results from evolutionary simulations on the relationship between recombination rate and *d* in models with selection in each of three simulated *N_e_* and three time points after the split from the ancestral pop_A_. The plots on the left are the same as Fig. 4, but additional dashed lines show mean *d* for each *N_e_* and time point from the neutral model (without gene conversion) to facilitate comparisons between models with and without selection, and with a mild (left panel) and a more severe bottleneck (right panel).

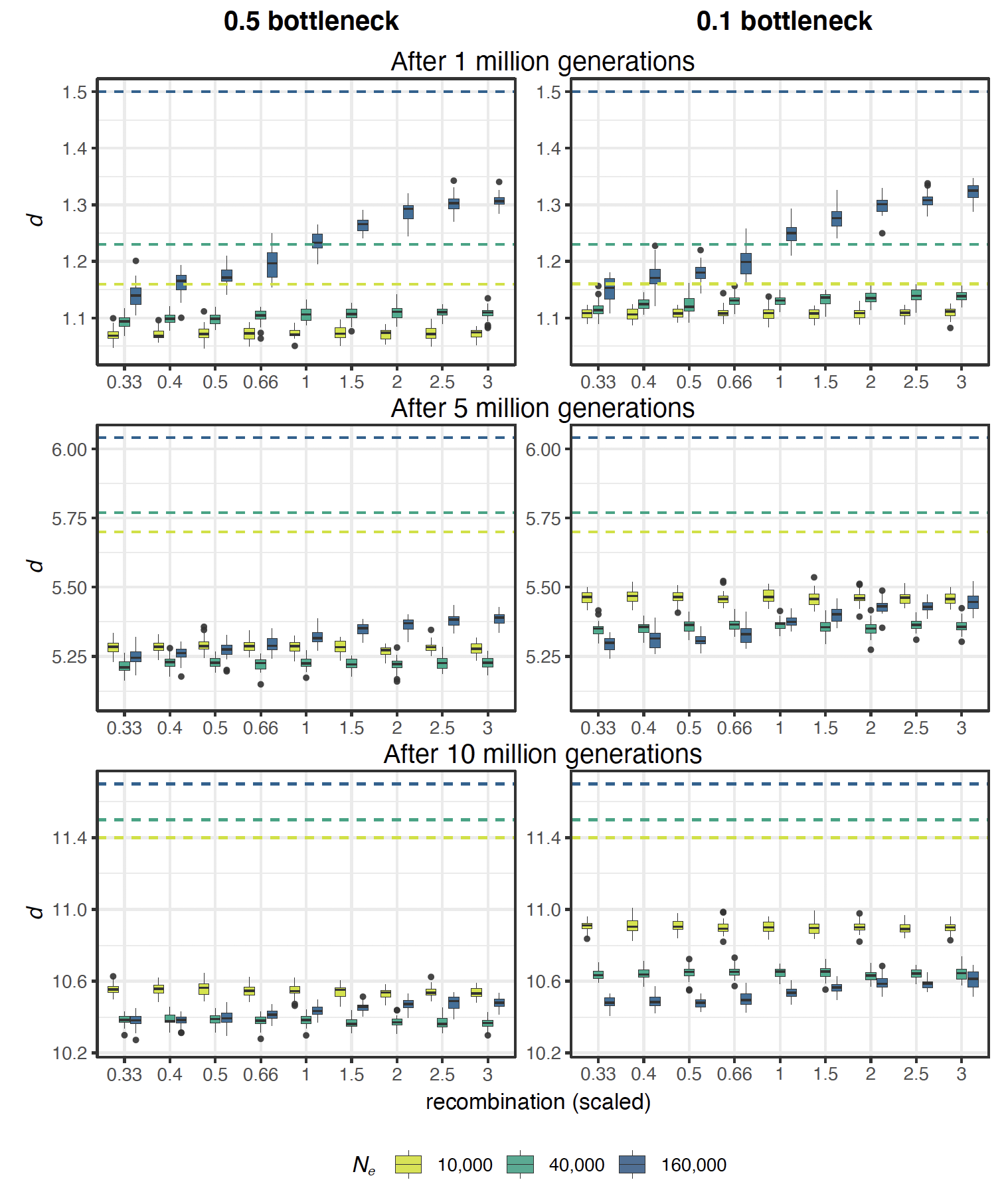

**Fig. S5.** Boxplots summarizing comparison between results obtained from simulations where only recombination varied and chromosome size was fixed (1 Mb – gray in the plot) and simulations where chromosome size varied according to recombination rate (3 Mb, 0.66 Mb, and 0.33 Mb – yellow, light green, and dark green, respectively). The plots show the results for π in the ancestral population right before the split based on a model with selection and no gene conversion. The different columns (associated with increasingly darker shades) show results for of the three simulated *N_e_*, and the different row show results for different regions of the chromosome (all, only genes, only intergenic areas).
